# Gli1^+^ mesenchymal progenitors contribute to multilineage differentiation in a mouse model of post-traumatic joint injury

**DOI:** 10.1101/2022.06.11.495765

**Authors:** Jenny Magallanes, Nancy Q Liu, Jiankang Zhang, Yuxin Ouyang, Tadiwanashe Mkaratigwa, Fangzhou Bian, Ben Van Handel, Tautis Skorka, Frank A. Petrigliano, Denis Evseenko

## Abstract

Complex injury and open reconstructive surgeries of the knee often lead to joint dysfunction that may alter the normal biomechanics of the joint. Two major complications that often arise are excessive deposition of fibrotic tissue and acquired heterotopic endochondral ossification. Knee arthrofibrosis is a fibrotic joint disorder where aberrant buildup of scar tissue and adhesions develop around the joint. Heterotopic ossification is ectopic bone formation around the periarticular tissues. Even though arthrofibrosis and heterotopic ossification pose an immense clinical problem, limited studies focus on their cellular and molecular mechanisms. Effective cell-targeted therapeutics are needed, but the cellular origin of both knee disorders remains elusive. Moreover, all the current animal models of knee arthrofibrosis and stiffness are developed in rats and rabbits, limiting genetic experiments that would allow us to explore the contribution of specific cellular targets to these knee pathologies. Here, we present a novel mouse model of post-traumatic joint injury where surgically induced injury and hyperextension of the knee lead to excessive deposition of disorganized collagen in the meniscus, synovium, and joint capsule in addition to formation of extra-skeletal bone in muscle and soft tissues within the joint capsule. As a functional outcome, arthrofibrosis and heterotopic endochondral ossification coupled with a significant increase in total joint stiffness were observed. By employing this model and genetic lineage tracing, we also demonstrate that Gli1^+^ mesenchymal progenitors proliferate after joint injury and contribute to fibrotic cells in the synovium and ectopic osteoblasts within the joint capsule. These findings demonstrate that Gli1^+^ cells are a major cellular contributor to knee arthrofibrosis and heterotopic ossification that manifests after knee injury. Our data collectively shows that genetic manipulation of Gli1^+^ cells in mice may offer a platform for identification of novel therapeutic targets to prevent chronic knee joint dysfunction after chronic injury.

## INTRODUCTION

Arthrofibrosis and acquired heterotopic endochondral ossification are two recognized sequelae of complex knee injury and orthopaedic surgeries of the knee. They can be initiated by different triggers such as fractures, traumatic events, acute trauma, and combat related injuries. Moreover, although knee surgeries, such as total knee arthroplasty (TKA), are often reliable surgeries that relieve conditions of the knee, arthrofibrosis and acquired heterotopic endochondral ossification continue to compromise the outcome of many patients (Abdel et al., 2012). Knee arthrofibrosis is characterized by the excessive buildup of scar tissue and adhesions in the joint, while heterotopic endochondral ossification is defined by the aberrant formation of masses of bone through a cartilage intermediate in extra-skeletal soft tissues. Both post-traumatic joint traumas can cause knee stiffness and restrict the range of motion of the affected joint. (Cheuy et al., 2017; Pagani et al., 2021). Clinically, patients present with limited knee range of motion and chronic pain with routine daily activities. In several cases, these post-traumatic joint injuries may become progressive with fibrous tissue and/or ectopic bone thickening and tightening the entire capsule, causing other health problems, including nerve compression and pressure ulcers (McCarthy and Sundaram, 2005). Once arthrofibrosis or heterotopic ossification (HO) develops, surgical removal is the only effective treatment (Cheuy et al., 2017; Mundy et al., 2021).

These post-traumatic joint complications could be prevented by interventions that modulate their cellular cascades; yet their fundamental mechanisms are not well-understood. One limiting factor has been the lack of relevant animal models. Appropriate animal models for post-traumatic joint injuries are key to evaluate new treatments. Mouse models are essential because they enable the use of genetically modified mice to investigate the pathogenesis of joint contractures. Although reproducible knee arthrofibrosis and HO models have been generated in a variety of species, such as rat and rabbit (Baranowski et al., 2018; Hazlewood et al., 2018; Kaneguchi et al., 2021; Steplewski et al., 2021), there are no reliable mouse models that recapitulate the important features of knee arthrofibrosis or acquired HO observed in humans and larger animal models post-chronic injury. The only reproducible mouse models of HO are of genetic HO, which require genetic modifications leading to enhanced BMP signaling (Kan et al., 2019). But in human patients, the rare genetic causes of HO have a different clinical presentation and severity than the far more common acquired cases. Current models of knee arthrofibrosis and acquired HO involve immobilizing the knee joint by external fixation using a splint or joint bandaging (Kan and Kessler, 2011; Tokuda et al., 2022). However, the pathophysiology observed in immobilization mouse models does not represent severe arthrofibrosis or HO as seen in patients that have undergone severe knee joint trauma post-surgery or post-injury. Post-traumatic joint injury is more complex and involves both hard and soft tissues in the knee joint as well as substantial bleeding into the synovial cavity.

The correct programming of mesenchymal progenitors is critical to coordinate normal wound healing after chronic injury (Julien et al., 2021; Whelan et al., 2020). However, during severe trauma, improper mesenchymal progenitor differentiation can be maladaptive, causing pathological healing. This altered programming of mesenchymal progenitors is manifested in the process of arthrofibrosis and acquired HO (Huang et al., 2021; Usher et al., 2019). Despite the eminent therapeutic potential of this cell type, there is a lack of *in vivo* studies that demonstrate its contribution to these post-traumatic knee joint pathologies. Previous studies have shown that perivascular Gli1^+^ cells from multiple organs (kidney, lung, liver, and heart) in addition to Gli1^+^ cells from the mouse incisor express typical markers of mesenchymal progenitors and have trilineage differentiation capacity toward chondrocytes, adipocytes, and osteoblasts *in vitro* (Kramann et al., 2015; Shi et al., 2017; Zhao et al., 2014). *In vivo* studies have demonstrated that Gli1^+^ cells give rise to multiple cell types associated with the skeleton in the long and craniofacial bones (Shi et al., 2017; Zhao et al., 2015). Moreover, in pathological conditions, Gli1^+^ cells can also proliferate after kidney, lung, liver, or heart injury to generate fibrosis-driving myofibroblasts (Kramann et al., 2015). Finally, Gli1^+^ cells have also been identified as a major cellular origin of genetic HO (Kan et al., 2018; Kan et al., 2019). These findings provide a rationale for potentially targeting Gli1^+^ cells to mitigate their contribution to arthrofibrosis and acquired HO.

In this study, we have developed a surgically induced mouse model of post-traumatic knee joint injury. We have taken some elements of models developed in rats and rabbits to generate a mouse model. This model not only reproducibly results in knee arthrofibrosis, but also in acquired HO (not genetic). We utilized this injury model in Gli1-CreER^+/ERT2^; tdTomato^+/-^ mice to identify, lineage trace, and analyze the fate of Gli1^+^ cells post-injury. Using lineage tracing analyses, we found that the tdTomato^+^ population greatly contributes to joint fibrosis and acquired HO following injury. Our results show that these cells can differentiate into fibrotic and osteogenic lineages. Taken together, our findings demonstrate that we have developed a surgically induced mouse model of knee arthrofibrosis and acquired HO. We also show that Gli1 lineage cells are a unique cell population of mesenchymal progenitors that can undergo an aberrant fate change post-injury and are therefore a relevant therapeutic target.

## MATERIALS AND METHODS

### Mouse Lines and Breeding

All procedures involving animals were approved by the Institutional Animal Care and Use Committee (IACUC) of USC. This study was compliant with all relevant ethical regulations regarding animal research. We used the following mouse lines: Gli1-Cre^ERT2^ (JAX; 007913) and Rosa26tdTomato (JAX; 007914). Gli1-Cre^ERT2^ males were crossed to Rosa26tdTomato females to generate Gli1Cre^+/ERT2^;Rosa26tdTomato^+/-^ mice. All animals were on a C57BL/6 background. Genotyping was performed according to protocols provided by JAX. For lineage tracing studies, 7-to 10-week-old mice received tamoxifen (100 µg per gram of body weight) in corn oil via intraperitoneal injection twice 7 days before surgery.

### Surgical Procedures

In this IACUC-approved study, surgeries were performed on the left knee joint under sterile conditions and general anesthesia. Mice (18-22g) were anesthetized using isoflurane at 2% with oxygen carrier via nasal inhalation using a nose cone. Ophthalmic lubricant was applied to the eyes to prevent corneal drying while under anesthesia. Animals then received pre-operative doses of non-steroidal anti-inflammatory drug (NSAID) (5 mg/kg Carprofen) and Buprenorphine SR-LAB (1 mg/kg) subcutaneously. The surgical area was shaved with an electric clipper and prepped with 10% Povidone-Iodine and 70% Isopropyl Alcohol. Medial para-patellar arthrotomy was performed as previously described (Eltawil et al., 2009) under a dissection microscope (Leica) by inserting a microsurgical scalpel medially and proximally to the insertion of the patellar tendon on the tibia and extending it proximally until the attachment of the quadriceps muscle. The medial margin of the quadriceps was separated from the muscles of the medial compartment. The joint was then extended, and the patella dislocated laterally. The joint was then fully flexed to expose the femoral condyle. The condyle defect, patella suture, and joint hyperextension were then carried out (**Figure 1**). Finally, the wound site was carefully rinsed with sterile saline before closing the joint capsule with a 7-0 Vicryl suture (Ethicon, J488G) and skin with a 5-0 Vicryl suture (Ethicon, J385H). Following wound closure, animals were allowed free cage activity. Carprofen was injected subcutaneously for the next two days. The right knee joint served as uninjured controls.

**Figure 1:**
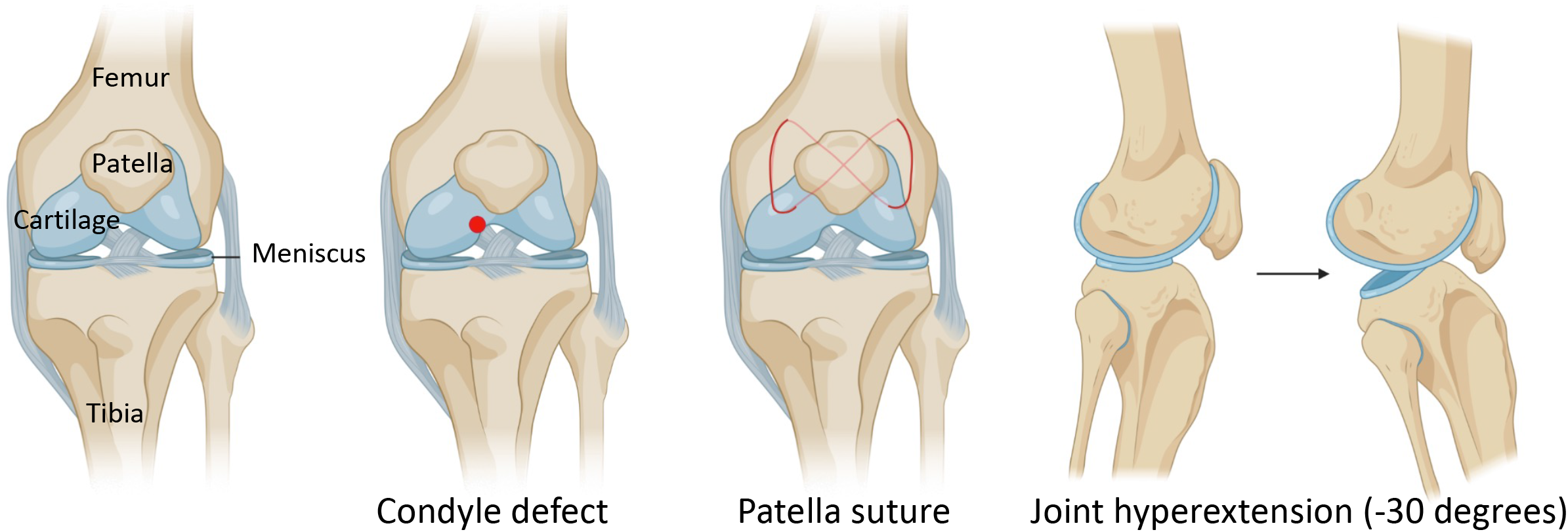
Schematic with surgical details of a mouse model of post-traumatic joint injury. The surgery consists of making a condyle defect in the femoral condyle, suturing the patella to dislocate it, and hyperextending the knee joint to disrupt the posterior joint capsule.

### Histology and Immunohistochemistry

Limb tissues were dissected and fixed in 10% formalin overnight. They were then decalcified with 14% EDTA, pH7.4, for 2 weeks at 4 °C. Decalcified tissue was then embedded in paraffin and cut at a thickness of 5 μm. Paraffin sections were deparaffinized and rehydrated by passage through xylene and 100, 95, and 70% ethanol. Tissue was then permeabilized with 0.5% PBST (0.5% Triton X-100 in PBS) for 20 min. Antigen retrieval was carried out in citrate buffer (pH 6.0) for 20 min at 95 °C.

For chromogenic labeling, inhibition of endogenous peroxidase activity was performed using 3% H_2_O_2_ for 10 min. Sections were blocked with 5% normal horse serum (NHS) in 0.5% PBST for 20 min and incubated overnight with primary antibody in 5% NHS in 0.5% PBST at 4 °C. Slides were then incubated at room temperature (RT) for 20 min in secondary antibody-HRP (Vector Laboratories, MP-7401). Finally, antibodies were visualized by peroxidase substrate kit DAB (Vector Laboratories, SK-4100). DAB-based staining was quantified by selecting all the pixels containing “brown” within the joint capsule in Photoshop. The total number of pixels identified were then counted using the histogram function.

For fluorescent labeling, sections were blocked with 5% normal donkey serum (NDS) in 0.5% PBST for an hour at RT. Then slides were incubated overnight with primary antibody in 5% NDS in 0.5% PBST at 4 °C. Slides were incubated at RT for an hour in secondary antibody and mounted with Fluoro-Gel II containing DAPI (Electron Microscopy Sciences, 17985). Slides were viewed using a Zeiss Axio Imager.A2 Microscope and images were taken using Axiocam 105 color (chromogenic detection) and Axiocam 105 (fluorescent detection) cameras with Zen 2 program. Standard microscope camera settings were used. Primary antibodies included the following: mCherry which also detects tdTomato (Novus Biologicals, NBP2-25158; 1:2000), αSMA (Thermo-Fisher Scientific, 701457; 1:100), and Collagen I (Thermo-Fisher Scientific, PA1-26204; 1:100). The secondary antibodies for immunofluorescence include donkey anti-rabbit Alexa Fluor 488 and donkey anti-chicken Alexa Fluor 594 (Thermo-Fisher Scientific; 1:1000).

Hematoxylin and eosin (H&E), Safranin O/Fast Green (SO/FG), and Picrosirius red staining were performed according to routine protocols. H&E staining was used to perform synovitis scoring as described (Krenn et al., 2006). SO/FG staining was used to perform Osteoarthritis Research Society International (OARSI) scoring as described (Pritzker et al., 2006).

### Range of motion assay

To assess joint contracture, knee range of motion was measured post-mortem six weeks after surgery. Measurements were taken 2 hours within euthanizing the mice. The skin was removed so that extension measurements were only composed of muscular and articular/capsular parts. Animals were then placed on their backs on a custom apparatus designed to be operated on a flat table. The femur was gripped by a serrated alligator clip that is drilled to an acrylic board. A cord was attached to the ankle joint, while the other side of the cord was attached to a sensitive sensor Force Gauge (Model M7-012, Mark 10, USA). Then, a knee extension movement from 90° flexion to 30°, which stretches the knee joint, but does not disrupt soft tissue, was applied. The mechanical force required to extend the knee was determined with the sensor (**Figure 4A**). To evaluate the contribution of articular/capsular factors to joint contracture, knee flexor muscles were removed, and range of motion was measured again.

### Dynamic weight bearing

As an indicator of joint pain, limb weight bearing was assessed in mice before and after surgery. The weight borne by each limb was measured using a dynamic weight bearing (DWB) system (BioSeb). The DWB is a test that evaluates spontaneous pain in freely moving mice and is based on an instrumented-floor cage with a video acquisition system. Mice were weighed right before the assay. For each measurement, each mouse was individually placed in a plexiglass chamber with 1936 floor sensors detecting pressure and a camera that detects the posture of the mouse. Data obtained from the sensor array and video were matched to calculate the weight applied on each limb over the dynamic movement for a 5 min period. The weight placed on each separate paw (g) was measured. A mean value for the weight placed on each limb was calculated over the entire testing period. Data were presented as a percentage of weight placed on the left/right hind paw relative to body weight: (weight on left/right limb / body weight) x 100.

### Micro-computed tomography (μCT)

Mouse limbs were harvested, fixed, and imaged 10-weeks post-injury with a μCT 50 cabinet microCT scanner (Scanco Medical). The following parameters were used to assess heterotopic ossification: 30μm resolution, 70kV energy, and 114 μA current. Visualization of 3D morphological structures was performed using Amira 3D 2021.1. software (Thermo-Fisher Scientific).

### Statistical Analysis

The number of biological replicates and types of statistical analyses for each experiment are indicated in the figure legends. All statistical analyses were performed using Prism (version 9.3.1, GraphPad Software Inc.). All results are presented as mean ± SEM where *p* values were calculated using two-tailed Student’s *t* test.

## RESULTS

### A mouse model of post-traumatic joint injury

As knee arthrofibrosis and acquired HO models have been generated in larger animal models, we sought to establish a similar model in the mouse knee joint. The surgical procedure involves drilling a hole of 0.9-mm in diameter with a Kirshner wire (K-wire) into the non-cartilaginous part of the medial distal femoral condyle close to the attachment site of the posterior cruciate ligament (PCL) to create a full thickness cartilage injury. This lesion mimics a juxta-articular fracture and causes bleeding into the joint (Hildebrand et al., 2004; Lake et al., 2016). Next, a 5-0 absorbable suture was used underneath the patella to dislocate it and increase friction in the trochlear grove. Finally, we used a hyperextension of -30° of the joint to disrupt the posterior joint capsule as previously described in larger animal models (Baranowski et al., 2018; Li et al., 2013; Sun et al., 2016). The knee joint angle is defined as the angle between the longitudinal axis of the femur and the longitudinal axis of the lower leg (**Figure 1**).

We next addressed the effect of our surgical model on joint morphology by performing histology following injury. Our data demonstrates that this procedure is effective in producing arthrofibrosis 6 weeks post-surgery as visualized by higher levels of fibrosis development in operated knees compared to control uninjured knees. The knee joints of mice that underwent surgery displayed higher levels of picrosirius red, showing high collagen deposition within the synovium, joint capsule, and meniscus (**Figure 2A**). When tissues within the joint capsule, including the synovium, are affected by fibrosis, they become thicker. Therefore, we assessed joint capsule thickness by measuring its width, which includes all tissues from the surface of the skin to the synovium lining. Capsular thickness was significantly increased (p=0.0003) in the surgical group in comparison with the uninjured group (**Figure 2B**). Operated mice also expressed higher levels of notorious pro-fibrotic markers: α-SMA and Collagen I (**Figure 2C**). Chondrogenesis is an essential step of endochondral HO. By 6 weeks, ectopic Safranin O staining was observed within the joint capsule (**Figure 2D**). Safranin O/Fast Green (SO/FG) staining confirmed cartilage formation with chondrocytes flanking other areas that had developing marrow space or undergone mineralization (**Figure 2D**). These data indicate that our novel post-traumatic injury model can produce both knee arthrofibrosis and developing HO by 6 weeks post-surgery.

**Figure 2:**
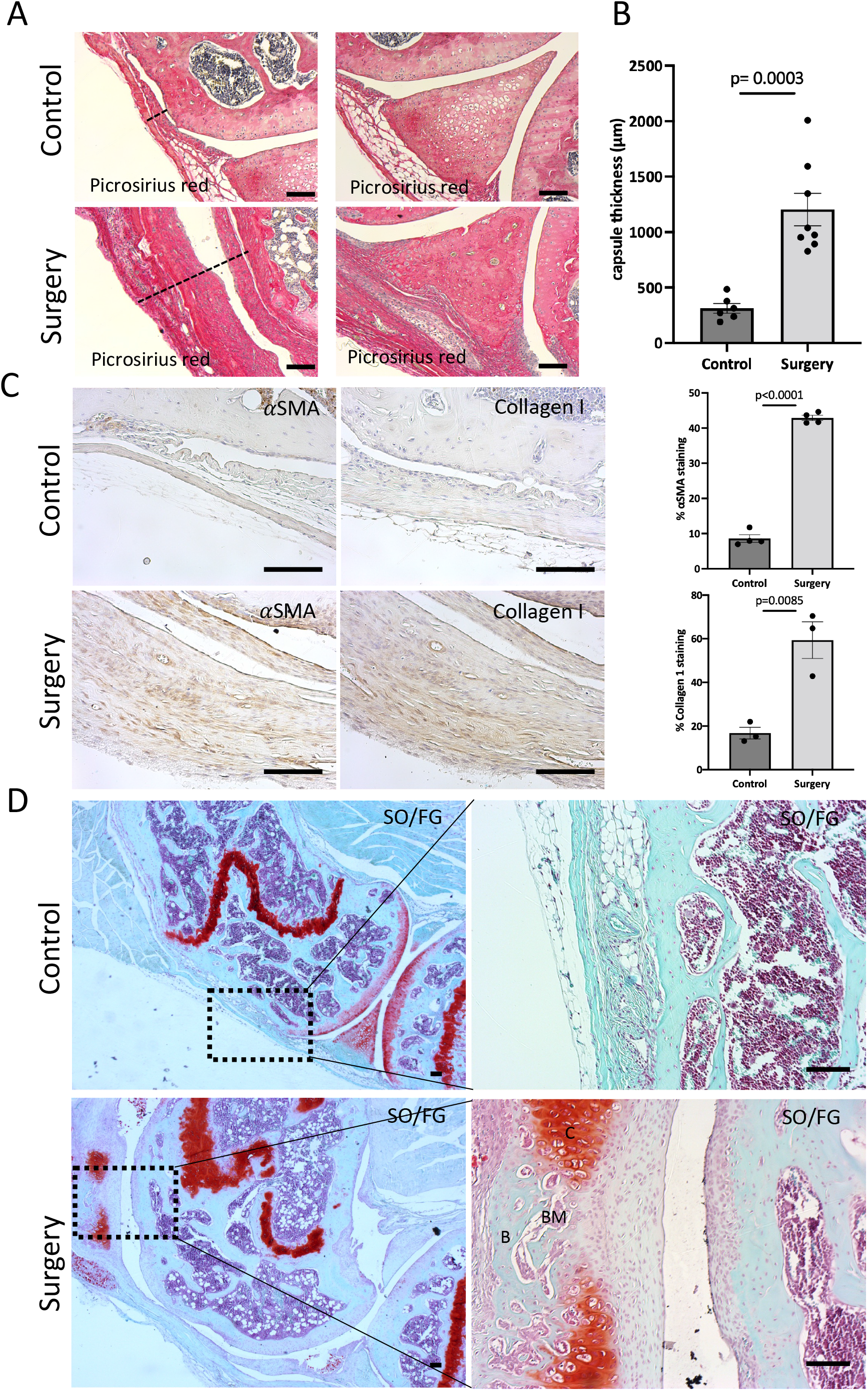
Surgery results in knee arthrofibrosis and acquired heterotopic ossification. **(A)** Histological staining of operated knee joints 6 weeks post-injury shows higher collagen deposition in the joint capsule and meniscus when compared to uninjured controls. Picrosirius Red staining delineates collagen deposition (red). **(B)** Surgery increased capsule thickness, including synovial hyperplasia **(C)** Immunohistochemistry and quantification of α-SMA and Collagen I. Both were highly expressed in operated mouse joints. **(D)** Safranin O/Fast Green (SO/FG) shows progression of endochondral HO in operated joints. B, bone; BM, bone marrow; C, cartilage. In all panels, scale bars = 100um; n= 3-8. *p* values were calculated using two-tailed Student’s *t* test. All error bars represent mean ± SEM.

Synovitis associated with joint injury can promote synovial angiogenesis, which in turn accelerates inflammation and leads to synovial fibrosis (Zhang et al., 2021). At six weeks post-surgery, morphological signs of synovitis were prominent in knee joints that underwent surgery. Operated knee joints showed significant synovitis (p<0.0001) when analyzing H&E staining using synovitis scoring system (Krenn et al., 2006) (**Figure 3A**). This score considers three components of synovitis: enlarged synovial lining cell layer (p<0.0001), increased resident cell density (p<0.0001), and enhanced inflammatory infiltration (p<0.0001) (**Figure 3A**). We next evaluated the structural integrity of the articular cartilage using SO/FG staining and OARSI scoring system (Pritzker et al., 2006). Six weeks after surgery, the articular cartilage in the operated knees exhibited significant (p=0.0012) pathological changes characterized by Safranin O loss, cartilage fibrillation, and cartilage erosion (**Figure 3B**). Our findings show that our injury model also leads to synovitis and cartilage degeneration, likely due to the severity of injury used in this model.

**Figure 3:**
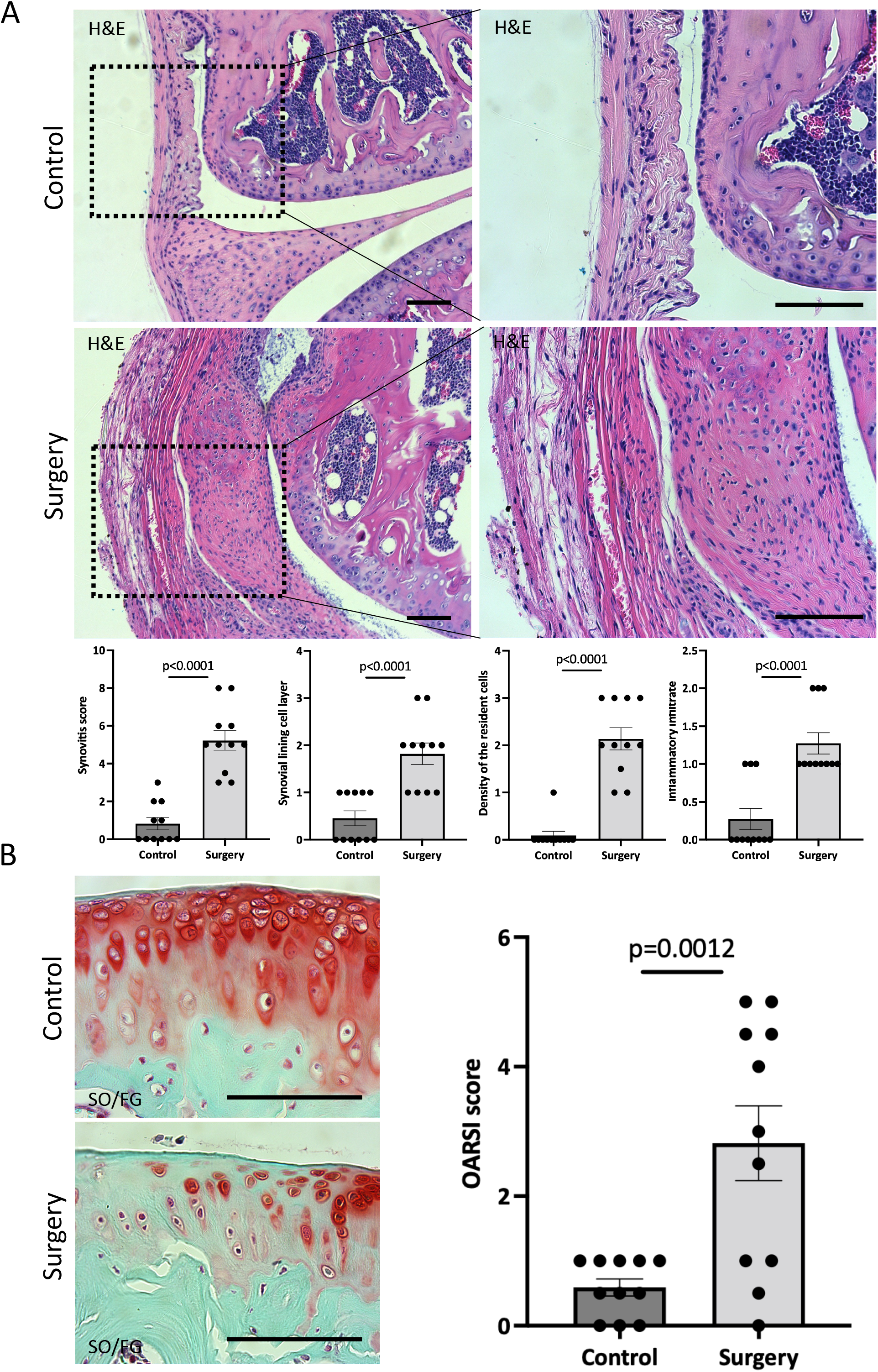
Knee joints exhibit synovitis and articular cartilage degeneration after surgery. **(A)** Quantitative assessment of synovitis in control and operated mouse knee joints 6 weeks after surgery. H&E staining shows that operated joints exhibit high-grade synovitis (p<0.0001). The scoring system assesses synovial lining cell layer, density of resident cells, and the inflammatory infiltrate. **(B)** Histological staining and quantitative assessment of cartilage degradation of mouse knee joints 6 weeks post-surgery. SO/FG delineates proteoglycans. OARSI scoring system was used to quantify the extend of cartilage damage. In all panels, scale bars = 100um; n=11. *p* values were calculated using two-tailed Student’s *t* test. All error bars represent mean ± SEM.

### Post-traumatic joint injury leads to loss of range of motion and pain

To measure the biomechanical effects of our post-traumatic joint injury model, joint stiffness was assessed. Knee stiffness following traumatic joint injury is a common clinical outcome of arthrofibrosis and HO. Our group previously developed a novel testing methodology to measure joint stiffness in rodent knees (Markolf et al., 2016) and has successfully adopted it to mouse studies. Mice were examined using a highly sensitive sensor force gauge to quantify the range of motion and contracture (**Figure 4A**). To assess that, we measured the mechanical force required to extend the knee from 90° flexion to 60°. The required force to extend the left or right knee before surgery showed no significant difference (p=0.1374), indicating that the right leg could be used as a control for the operated left leg (**Figure 4B**). However, post-surgery, the force required to extend the operated left leg was significantly higher (p<0.0001) than the force required to extend the right uninjured leg (**Figure 4C**). To evaluate the contribution of only articular/capsular factors to joint contracture, knee flexor muscles were removed, and range of motion was measured again. Our data shows that after removing the flexor muscles from the legs, the required force to extend the left or right knee before surgery showed no significant difference (p=0.1926) (**Figure 4D**), but it was significantly higher (p<0.0001) post-surgery (**Figure 4E**). These results show that operated knee joints have detectable joint stiffness by 6 weeks post-surgery, similar to the joint stiffness observed in human patients with knee arthrofibrosis or HO.

**Figure 4:**
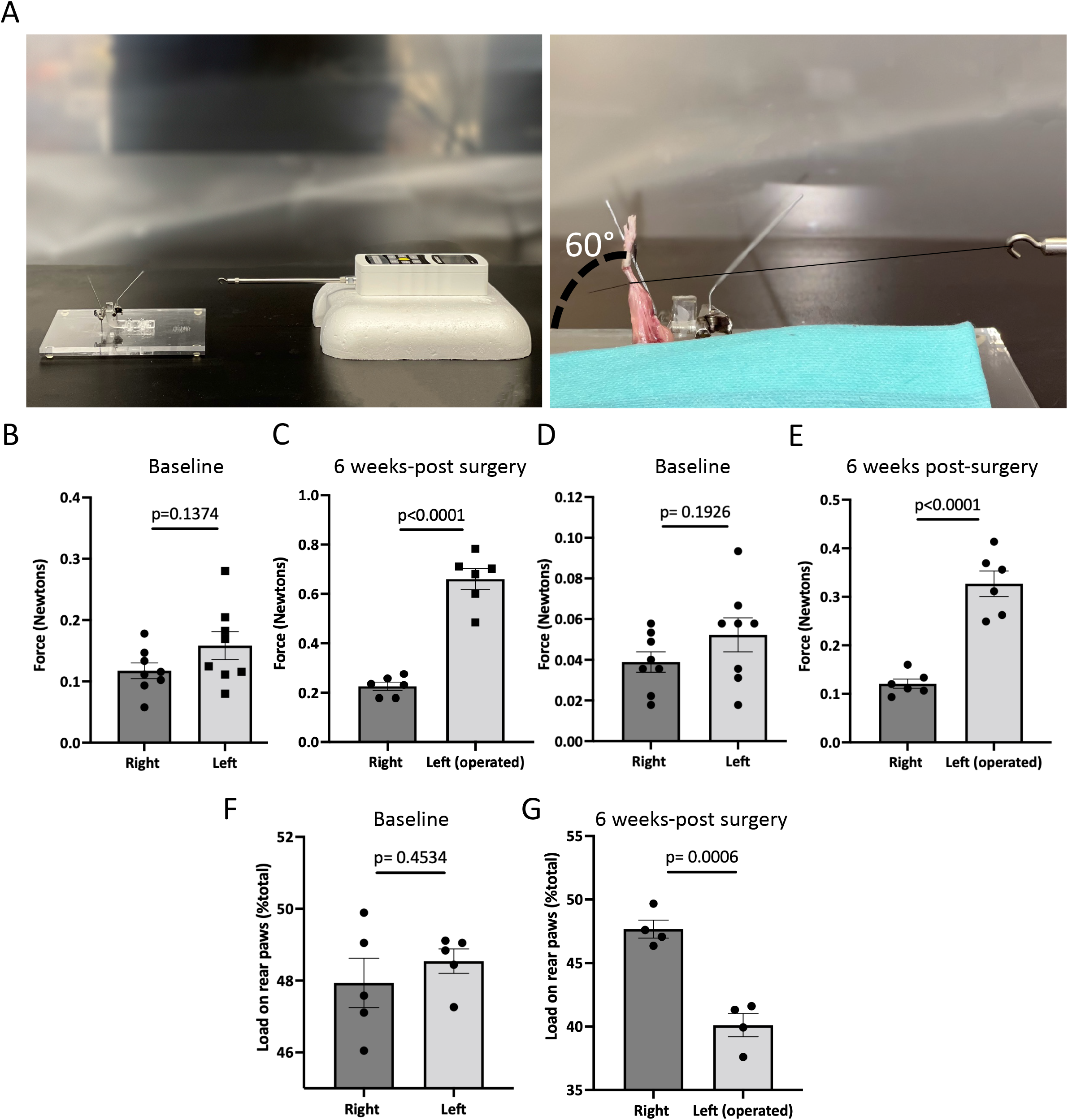
Post-traumatic joint injury leads to knee stiffness and pain. **(A)** Setup and apparatus for the knee range of motion assay. A thread was tied to the ankle. A force gauge apparatus was tied to the other end of the thread. Then the mechanical force required to extend the knee from 90° flexion to 60° was determined with the sensor. **(B)** Force required to extend the knee from 90° flexion to 60° before and **(C)** 6 weeks post-surgery. **(D)** Force required to extend the knee from 90° flexion to 60° before and **(E)** 6 weeks post-surgery after knee flexor muscles were removed. **(F)** Percentage of weight placed on the left rear paw vs right rear paw before **(G)** and after surgery. *P* values were calculated using two-tailed Student’s *t* test. All error bars represent mean ± SEM.

To assess joint-related pain, we used dynamic weight bearing (DWB) to evaluate the ability to place weight on the operated limb. There was no significant difference (p=0.4534) between the amount of weight placed on the left vs right rear limb before surgery (**Figure 4F**). However, mice put significantly less (p=0.0006) weight on the left operated limb compared to uninjured contra-lateral controls (**Figure 4G**). This shows that weight bearing is affected in those animals that underwent surgery, indicating persistent pain behavior. These measures of functional status are important for more accurate assessment of novel pharmacological therapeutics.

### Gli1^+^ cells contribute to knee arthrofibrosis

We next asked whether resident mesenchymal progenitors are activated post-injury and contribute to knee arthrofibrosis. Since Gli1 has been identified as a faithful marker for fibrosis-driving mesenchymal progenitors in solid organs, we sought to characterize Gli1^+^ cells in the joint post-injury. Gli1Cre^+/ERT2^ driver mice were crossed to a tdTomato reporter for inducible genetic labeling. To trace the fate of Gli1^+^ cells, Gli1Cre^+/ERT2^;tdTomato^+/-^ mice received tamoxifen for 2 days to induce Gli1-specific expression of tdTomato (**Figure 5A**). Mice were then subjected to our post-traumatic joint injury model 7 days after the last tamoxifen dose to eliminate any possibility of Cre recombination after injury. Finally, mice were allowed to move freely until they were euthanized 6-weeks post-surgery (**Figure 5A**). Lineage tracing of Gli1^+^ cells labeled prior to surgery revealed that these cells undergo massive expansion by 6 weeks post-surgery and significantly contribute to arthrofibrosis, including thickening of the synovial capsule (**Figure 5B**). Imaging further demonstrated that Gli1^+^ cells also acquire expression of α-SMA, indicating myofibroblast differentiation at the molecular level (**Figure 5C**). These results show that Gli1^+^ cells get activated, expand, and differentiate into fibrotic cells following surgically induced arthrofibrosis.

**Figure 5:**
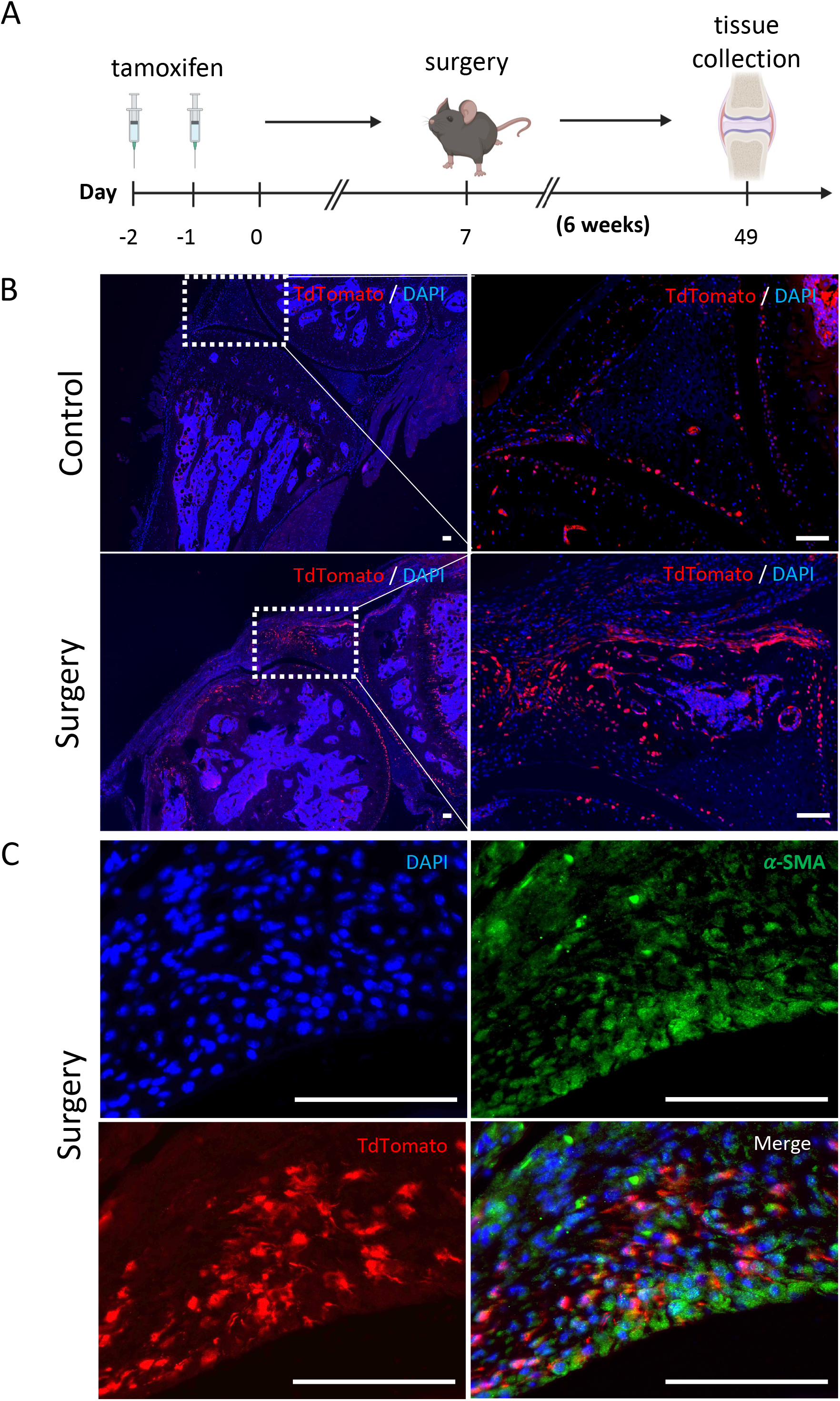
After knee injury, Gli1^+^ cells contribute to arthrofibrosis. **(A)** Experimental design: Gli1-CreER^+/ERT2^; tdTomato^+/-^ mice were injected with tamoxifen twice, surgery was carried out 7 days after the last tamoxifen dose. Mice were sacrificed 6 weeks post-surgery. **(B)** Gli1^+^ cells contribute to the pool of fibrotic cells post-injury as can be observed by immunofluorescence (IF) for the tdTomato protein (red). Representative images of knee joints of the control and surgical group 6 weeks post-surgery. **(C)** Operated knee joint co-stained with tdTomato and α-SMA, indicating that Gli1+ cells can differentiate into myofibroblasts. In all panels, scale bars = 100um.

### Gli1^+^ cells also contribute to acquired heterotopic endochondral ossification

As previously mentioned, chondrogenesis is a necessary step of endochondral HO. We sought to determine whether the bone islands that we previously found within the joint capsule of operated knees (**Figure 2D**) also originated from Gli1^+^ cells. Lineage tracing showed that Gli1^+^ cells contributed significantly to HO as we observed tdTomato expression in both chondrocyte as well as osteoblast-like cells (**Figure 6A**). To ensure that the observed morphology of what we believed would eventually completely ossify and become HO was not an artifact, we also analyzed the legs via μCT 10 weeks post-surgery. Similarly to the 6-week timepoint, to trace the fate of Gli1^+^ cells, Gli1Cre^+/ERT2^;tdTomato^+/-^ mice received tamoxifen for 2 days to induce Gli1-specific expression of tdTomato. After seven days, mice were subjected to our post-traumatic joint injury model and were euthanized 10-weeks later (**Figure 6B**). We then conducted μCT imaging to characterize the presence and location of HO islands in the knee joint area. HO formation was observed in the operated joints in various periarticular regions with a predominant ectopic bone deposition in the frontal peri-patellar area (**Figure 6C**). These data indicate that Gli1^+^ cells contribute significantly to the pool of osteoblasts during HO formation and that these masses completely ossify 10 weeks post-surgery as seen in our μCT data.

**Figure 6:**
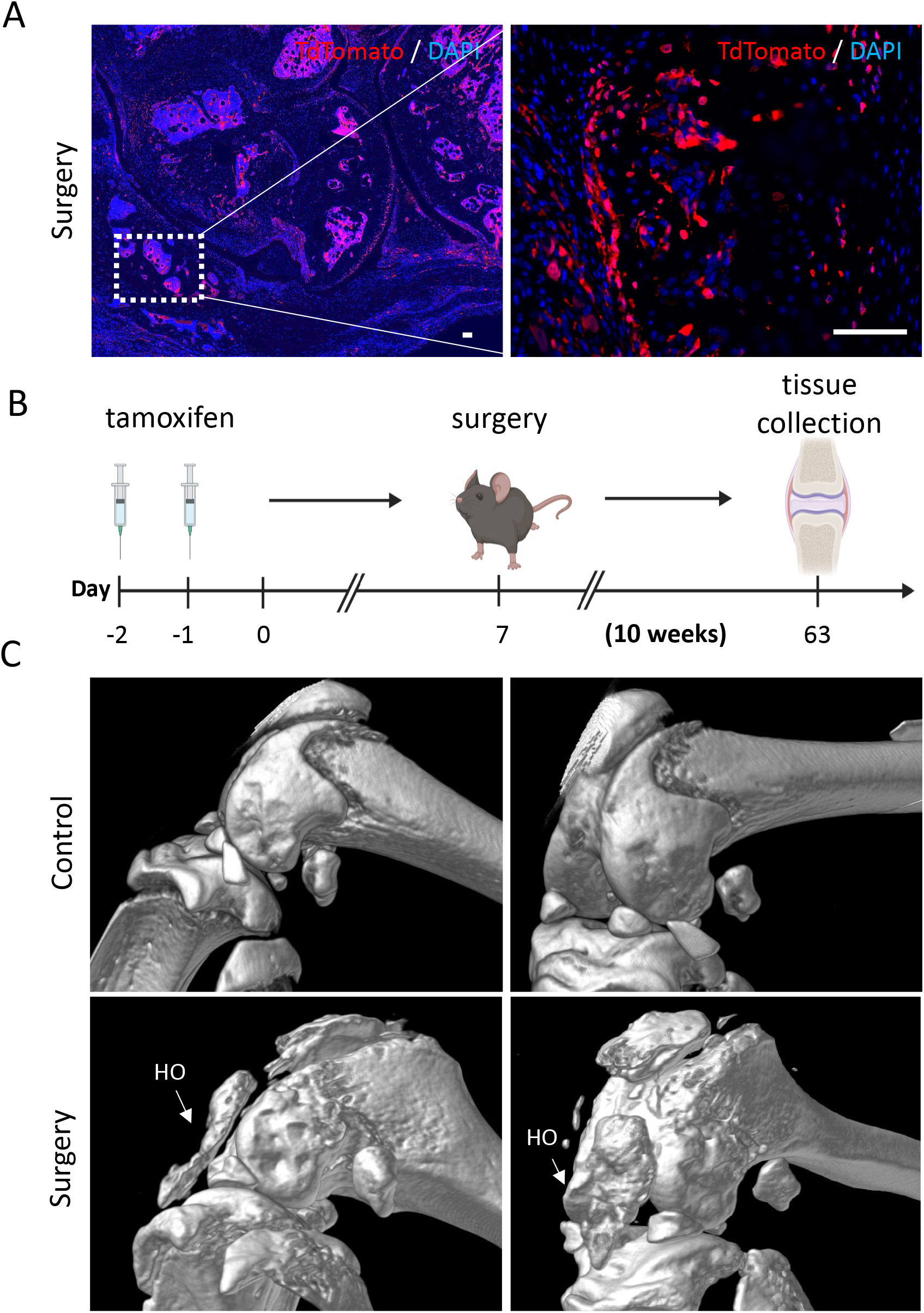
Gli1-expressing cells are also required for heterotopic ossification. **(A)** Gli1^+^ cells contribute to acquired heterotopic ossification. Representative images of ectopic bone in the knee joints of operated mice 6 weeks post-surgery. In all panels, scale bars = 100um. **(B)** Experimental design: Gli1-CreER^+/ERT2^; tdTomato^+/-^ mice were injected with tamoxifen twice, surgery was carried out at 7 days after the last tamoxifen dose. Mice were sacrificed 10 weeks post-surgery. **(C)** MicroCT analysis 10 weeks post-surgery shows HO formation in operated joints.

## DISCUSSION

To pursue the potential pathways involved in post-traumatic knee injuries, a sound animal model is necessary to understand the cellular and molecular processes leading to joint stiffness. Ideally, such models should reflect the clinical conditions involved to be able to identify potential targets for pharmacological intervention. In this study, we have described the development of a mouse model of post-traumatic joint injury. Micro-surgical techniques were developed for creating relevant tissue injuries representing chronic knee joint injuries that occur in humans. This surgical procedure successfully led to both knee arthrofibrosis and acquired HO uniformly present in all operated knee joints. Histologically, injured joints had capsule hyperplasia that expressed typical markers of myofibroblasts: α-SMA and Collagen I. Moreover, μCT analysis of the affected knees revealed ectopic bone, a feature of HO. 6-weeks post-surgery, operated joints also exhibited synovitis and cartilage degeneration.

We also demonstrated that our model led to restricted range of motion of the operated knee joints when compared to uninjured joints. Therefore, the mouse model developed in this study led to significantly altered joint mechanics that mimic symptoms of knee arthrofibrosis and HO in human patients. A novel approach was the use of DWB as a surrogate assessment of joint-related pain. The pattern of weight bearing of the rear limbs of operated knee joints clearly differed from the uninjured ones. This is consistent with the pain expected on operated limbs as seen in human patients that struggle with knee arthrofibrosis and/or HO. DWB, thus, offers a valuable non-invasive method for assessment of pain in mouse models of post-traumatic joint injury.

Various larger animal models of knee arthrofibrosis and acquired HO have been developed in the field. However, to our knowledge, there are no previous mouse models of surgically induced knee arthrofibrosis or acquired HO. Instead, previous studies have used joint immobilization models in mice to study the development of arthrofibrosis and acquired HO. For example, another group used surgical tape and an aluminum splint to immobilize the knee joint of mice for 4 weeks. This non-invasive method resulted in mild arthrofibrosis, with some thickening of the posterior capsule and some loss of range of motion (Kan and Kessler, 2011; Tokuda et al., 2022). However, although joint immobilization models do mimic some superficial elements of knee arthrofibrosis and acquired HO, these experimental conditions do not coincide with clinical observations in post-traumatic joint injury.

Currently, an unmet clinical need exists for novel therapies to treat or prevent the development of knee arthrofibrosis and acquired HO. Existing non-surgical treatments include physical therapy and the systemic administration of NSAIDs (Usher et al., 2019). Anti-inflammatories; however, do not halt arthrofibrosis or HO, they are only prescribed to treat the chronic pain associated with both (Monument et al., 2013). Local low-dose radiation is often given for HO prophylaxis (Mishra et al., 2011). This treatment is given right after joint injury and aims to target local osteogenic progenitors by reducing their functioning and viability. Clinically, this treatment is associated with lower incidence of HO, but it is not ideal, wholly effective, and secondary tumors can develop from radiation (Lee et al., 2016). Surgical treatments are also available for both knee arthrofibrosis and HO. Some surgical interventions for knee arthrofibrosis include arthroscopic lysis, debridement of extracellular matrix (ECM), and open surgery to remove ECM and release tendons and ligaments (Usher et al., 2019). The HO can also be removed by surgery (Meiners et al., 1997). However, surgeries are prone to complications and could potentially trigger another round of arthrofibrosis or HO if accompanied by extensive tissue damage or if the preexisting fibrotic tissue or HO masses are not resected completely (Pavey et al., 2015). In sum, although diverse clinical treatment options are available, there is an urgent need for safer and more effective approaches. Targeted pharmacological interventions may have the potential to alter the outcome of severe knee joint trauma and thereby prevent and treat joint contractures, but more mechanistic studies are needed to understand the molecular and cellular mechanisms of posttraumatic arthrofibrosis and HO.

Little is known about the *in vivo* role of mesenchymal progenitors in joint pathologies. Many populations of multipotent progenitors have been identified in long bones and their role during development has been extensively studied (Akiyama et al., 2005; Méndez-Ferrer et al., 2010; Zhou et al., 2014). As previously mentioned, Gli1^+^ cells can differentiate into important lineages during development, but they can also differentiate into pathological cell types. These findings highlight a predominant role for the perivascular niche in the scarring process. Here we demonstrate that Gli1^+^ cells undergo proliferative expansion after surgery and differentiate into myofibroblasts and osteoblasts *in vivo* and thus contribute to both knee arthrofibrosis and HO. Therefore, targeting Gli1^+^ cells might be beneficial in regenerative medicine by modifying the way these cells differentiate after injury. Understanding how these cells promote an abnormal wound healing response to injury could improve strategies to reverse or halt arthrofibrosis and HO.

In summary, this study presents a novel clinically relevant model of posttraumatic knee arthrofibrosis and HO in mice. Our data collectively shows that genetic manipulation of Gli1^+^ cells in mice may offer a platform for identification of novel therapeutic targets to prevent severe knee joint pathology after chronic injury. Finally, the establishment of this mouse model of post-traumatic joint injury provides an opportunity to test the roles of other cell populations in knee arthrofibrosis and acquired HO.

## Author Contributions

JM, NQL, JZ, YO, TM, FB, TS performed all experiments. JM, NQL, BV, and DE conceptualized the study and interpreted the data. JM wrote the manuscript. DE and FAP revised and approved the manuscript.

## Acknowledgments

The authors thank the Molecular Imaging Center core in the Department of Radiology, Keck School of Medicine, University of Southern California. All schematics were created with Biorender.com.

